# Mechanistic investigation of SARS-CoV-2 Omicron variant spike mutants via full quantum mechanical modeling

**DOI:** 10.1101/2021.12.01.470748

**Authors:** Marco Zaccaria, Luigi Genovese, Brigitte E. Lawhorn, William Dawson, Andrew S. Joyal, Jingqing Hu, Patrick Autissier, Takahito Nakajima, Welkin E. Johnson, Ismael Fofana, Michael Farzan, Babak Momeni

**Author notes:** These authors contributed equally to this work (Marco Zaccaria, Luigi Genovese).

## Abstract

*Ab initio* quantum mechanical models can characterize and predict intermolecular binding, but only recently have models including more than a few hundred atoms gained traction. Here, we simulate ∼13,000 atoms to predict and characterize binding of SARS-CoV-2 spike variants to the human receptor ACE2 (hACE2). We compare four spike variants in our analysis: Wuhan, Omicron, and two Omicron-based variants. To assess binding, we mechanistically characterize the energetic contribution of each amino acid involved, and predict the effect of select single point mutations. We validate our computational predictions experimentally by comparing binding efficacy of spike variants to cells expressing hACE2. We argue that this computational model, QM-CR, can identify mutations critical for intermolecular interactions and inform the engineering of high-specificity interactors.

**One-Sentence Summary:** *Ab initio* modeling can predict the strength of SARS-CoV-2 variants’ binding to human cell receptor.

## Background

To initiate entry to a host cell, virions bind to a target cell receptor. Mutations within the viral population explore the available chemical space to favor stronger binding to host receptors. If overall more fit, variants with such mutations can become dominant. Over the course of the SARS-CoV-2 pandemic, several emerging SARS-CoV-2 strains had modifications at the receptor binding domain (RBD) of the viral spike protein (S) [1–4]. The S protein binds the angiotensin converting enzyme 2 (hACE2) receptor to initiate SARS-CoV-2 infection. We aim to mechanistically understand how S interacts with hACE2 to predict which viral variants best exploit the chemical space, and how. Under the assumption of increased S-hACE2 binding strength as one of the drivers of viral adaptation to human hosts, we can identify which mutations pose the highest risk to characterize a future successful SARS-CoV-2 variant. Developing such predictive mechanistic models can facilitate preemptive engineering of pharmacological countermeasures to block viral evolutionary trajectories towards stronger binding. Prepared and guided by this insight, vaccines or treatments can nip dangerous lineages in the bud, before they become dominant.

Predictive models based on Quantum Mechanics calculations can characterize intermolecular interactions at high resolution. A full quantum mechanical environment can be simplified and made accessible by focusing the observables to the amino acid level, in terms of variation and characterization of their contribution to the overall binding energy between two proteins, without requiring prior knowledge and parameterizations. Here, we employ the S-hACE2 binding as a case study to illustrate what *ab initio* full QM models can provide in predicting intermolecular binding involving thousands of atoms.

We implement the full QM model using the Quantum Mechanics Complexity Reduction (QM-CR) formalized in a recent work [5]. Starting from a representative 3D model of the molecules as input [6], the BigDFT program [7] calculates the electronic density matrix based on Density Functional Theory to extract inter- and intra-molecular interactions. The strength of inter-residue interactions is quantified through the Fragment Bond Order (FBO) [8]. FBO is calculated using the electronic structure of the system, in proximity of a given residue. FBO identifies the residues involved in a chemical interaction, namely the amino acids of the counter-ligand that share a non-negligible bond—above a set threshold—with the ligand. In contrast to a purely geometrical indicator, like the residue-residue distance, FBO enables a non-empirical identification of steric hot-spot interactions. Once a chemical interaction is identified, we assign to each residue a specific value in the overall contribution to the intermolecular binding. These quantities are outputs of the BigDFT code and represent two terms: i) short range: attractive via chemical bond, which is non-zero only if the cross-fragment electronic clouds overlap; ii) long range: electrostatic attractive/repulsive, defined from the electron distributions of each fragment; long-range terms allow identification of off-interface relevant residues. Ultimately, QM-CR provides a mechanistic *ab initio* representation of a given ligand-counter ligand interaction as the final output. A contact map summarizing relevant interactions between the Wuhan S RBD and hACE2 is available in our previous work on QM-CR [5].

## Results

Throughout this work, we use the 6M0J entry in the RCSB database [9] as our starting crystallographic structures. We assign pH 7 to histidine protonation and other titratable residues via the PDBFixer tool, in OpenMM [10]. To generate the crystal structures with point mutations, we optimized the geometry through structure relaxations via the AMBER FF14SB force field [11], also available in the OpenMM package. This procedure aims at understanding the first order deviations in bindings implied by the variant, which is here characterized as the list of mutations relative to the wild-type structure. While such structures do not exhaust all possible conformations at a finite temperature, this formalization has already been successfully applied to the S-hACE2 binding, in a previous work [5].

### Compared to Wuhan S RBD, Omicron binds to hACE2 more strongly

We imposed the following mutations to represent the Omicron variant’s S RBD: K417N, N440K, G446S, S477N, T478K, E484A, Q493K, G496S, Q498R, N501Y, and Y505H. We opted to leave out G339D, S371L, S373P, and S375F, since they are far from the interface and not electrostatically active. QM-CR highlighted how the mutations present in the Omicron variant lead to a different interaction pattern with hACE2 compared to the original strain. We characterized the role of each mutation in the interaction rearrangement according to the hot spot regions they define, and according to their stabilizing power on and off the interface. Omicron is predicted to have a much stronger binding enthalpy than Wuhan (Fig 1, lower right panel). The interaction pattern of Omicron with hACE2 (Figs 1 a & b) highlights a comparable chemical/short range contribution to binding, but a higher long-range/electrostatic contribution which strongly favors Omicron over Wuhan, overall.

**Fig 1.**
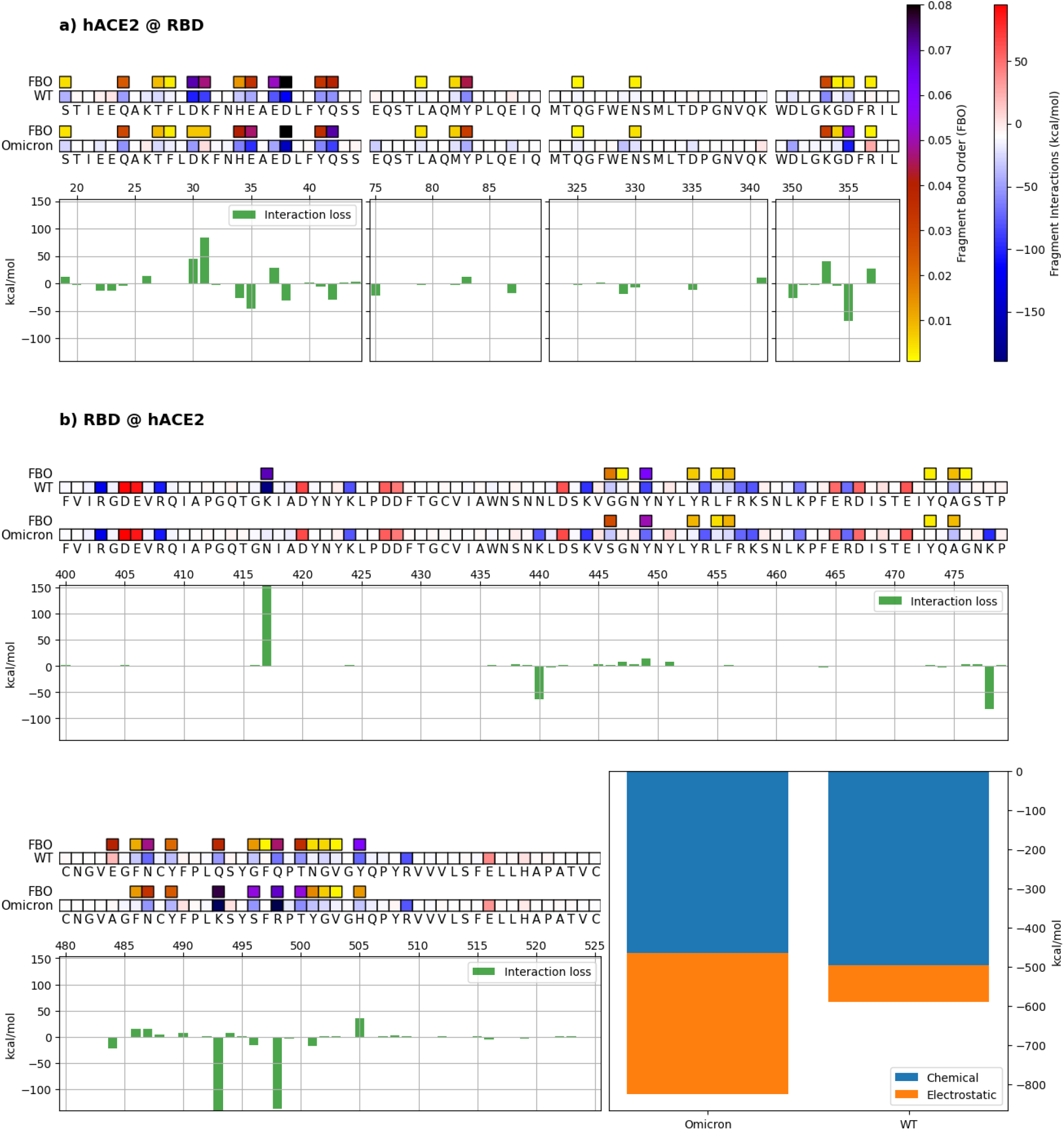
Mechanistic characterization of spike-hACE2 binding for the Wuhan (WT) and the Omicron variant spikes. Data are plotted on hACE2 (panel a) and the Wuhan (WT) and Omicron viral spikes (panel b). Amino acids are represented by the corresponding letters and numbered on each histogram’s horizontal axis. The contribution of each amino acid to the overall binding is indicated by red and blue squares for repulsive and attractive interactions, respectively. The energy scale is identical throughout all figures. Interface residues are highlighted by tiles on top, with darker colors indicating stronger interface values. Histograms underneath the sequences show the relative change in binding energy. Bottom right bars represent the overall binding energy of hACE2 with the Wuhan and Omicron variants, partitioned into chemical or electrostatic contributions.

### Some S RBD mutations can further strengthen its binding to hACE2

We tested whether QM-CR can predict the strength of hACE2 binding to hypothetical variants that may arise in the future. To this end, we examined two mutants that, at the start of our effort [12], had not yet emerged in epidemiological surveys: a) Omicron+L452R (now identified as Omicron BA.4/5), incorporating a mutation from the formerly dominant Delta variant; and b) Omicron+A484K, incorporating a mutation, from the once common Beta and Gamma variants, known to enhance hACE2 binding and antibody evasion [13]. QM-CR simulation predicted that L452R does not substantially alter the S-hACE2 binding strength; the increase in long-range contribution is canceled out by energy losses along the S structure. Conversely, QM-CR predicted that A484K dramatically increases the electrostatic component of the S-hACE2 binding, without compromising preexisting hot spots (Fig 2). In addition to the differences among the tested variants in overall binding energy, the interaction pattern of the Omicron+A484K includes new residues now participating in the contact interaction, such as L455, K484, and G502 on the RBD side.

**Fig 2.**
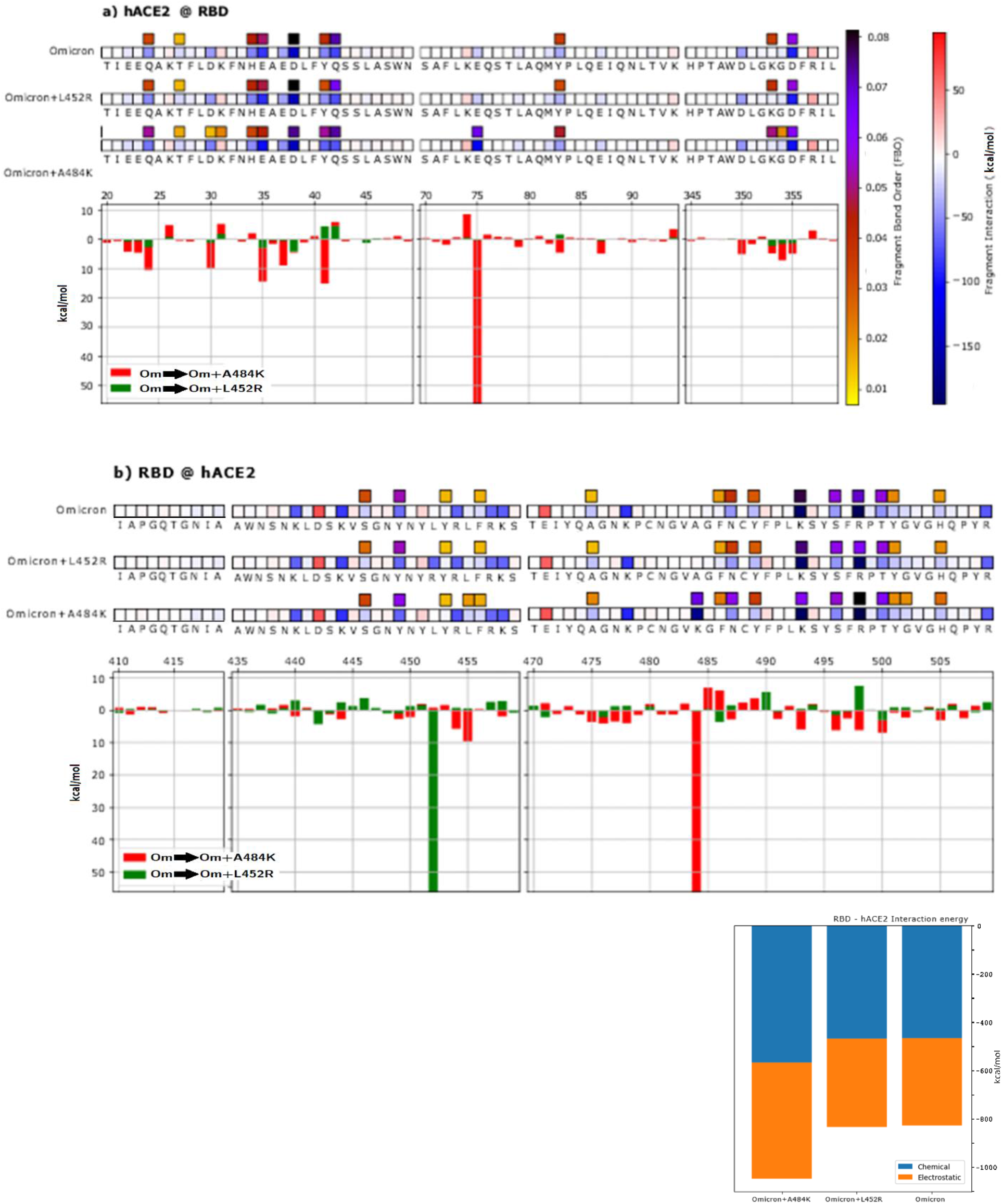
Characterization of the virtual mutations L452R and A484K of the Omicron variant. Data are plotted on the hACE2 primary structure (panel a), and on the Wuhan spike RBD (panel b), when binding to the Omicron, Omicron+L452R, or Omicron+A484K variants. The contribution of each amino acid to the overall binding is indicated by red and blue squares for repulsive and attractive interactions, respectively. Interface residues are highlighted by additional tiles on top, with darker colors indicating stronger interface values. Histograms underneath the sequences show the relative change in binding energy. Bottom right bars show the overall binding energy of hACE2 with the Wuhan and Omicron variants, partitioned into chemical or electrostatic contributions.

### Experiments validate the QM-CR predicted strength of S RBD binding to hACE2

To experimentally validate QM-CR predictions, we generated the RBD S-Fc fusion expression constructs of the Wuhan, Omicron, Omicron+A484K, and Omicron+L452R S proteins. The sequences were cloned into a plasmid expressing codon-optimized Wuhan S protein, the pCAGGS vector [14], using the NheI and SacI restriction sites. We tested the affinity of the novel variants by assessing their binding to 293T cells that express hACE2 on their surface. Using flow cytometry, we compared the strength of binding by assessing the mean fluorescence intensity (MFI) of cells bound to fluorescently stained S RBD (Fig 3).

**Fig 3.**
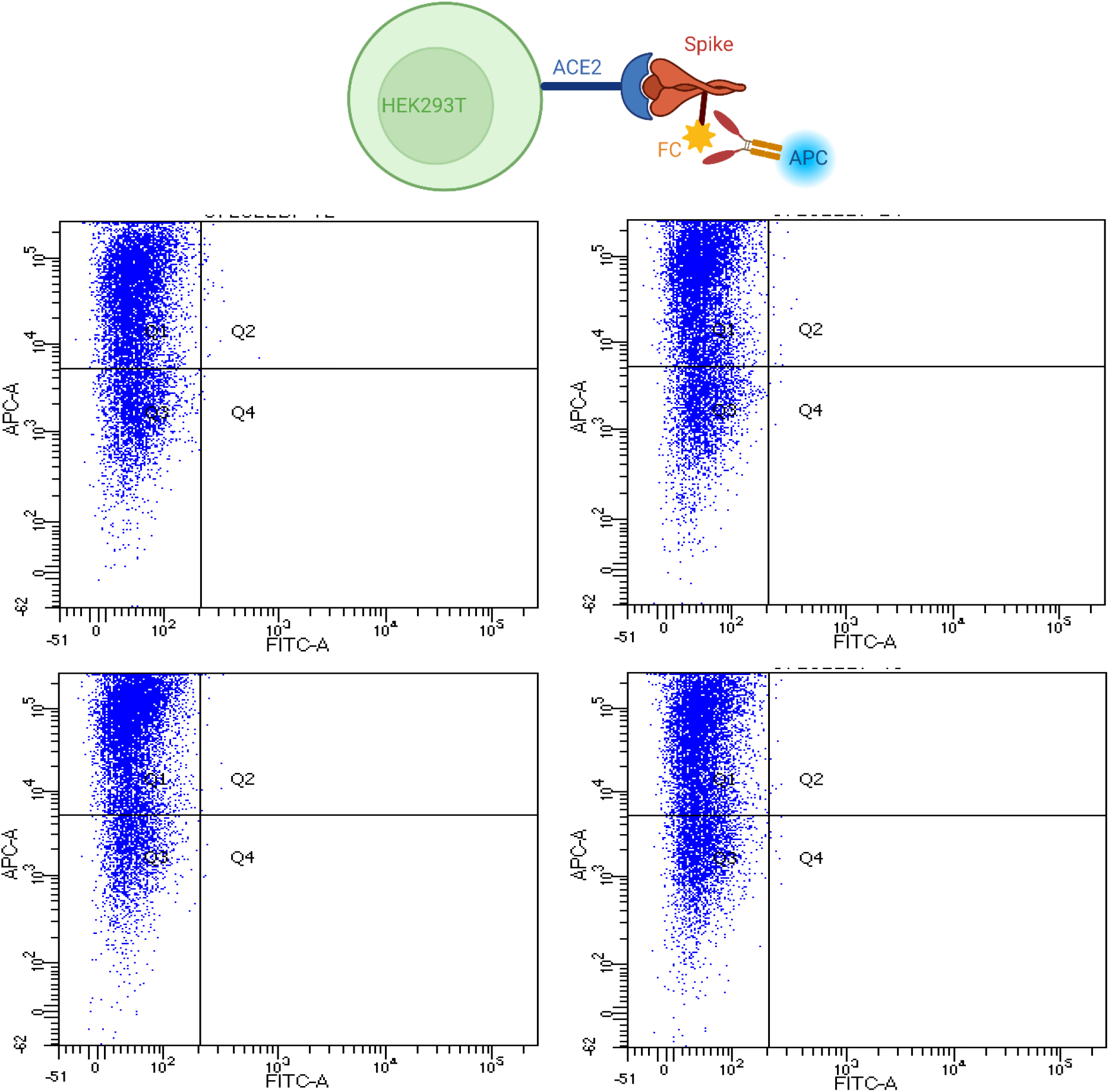
S-hACE2 Binding assays. For the binding experiment, the Spike protein’s FC tag is bound by an antibody carrying the APC fluorescent label. The four different S protein binding performances are expressed in mean fluorescence intensity (MFI) after binding.

In the experiment, Omicron+A484K displays the highest number of binding events, which correlates with QM-CR predictions (Fig 4). Omicron+L452R, Omicron, and Wuhan all follow the predicted trend (Fig 4). All pairwise comparisons of flow cytometry data for different spike variants showed statistically significant differences. Details of the *p*-values for flow cytometry data comparisons can be found in the supplementary information (Table S2). Fluorescence background values are available in the Supplementary Information: overall, the relevant quarter of the distribution is populated with less than 0.2% data points in the absence of the secondary antibody.

**Fig 4.**
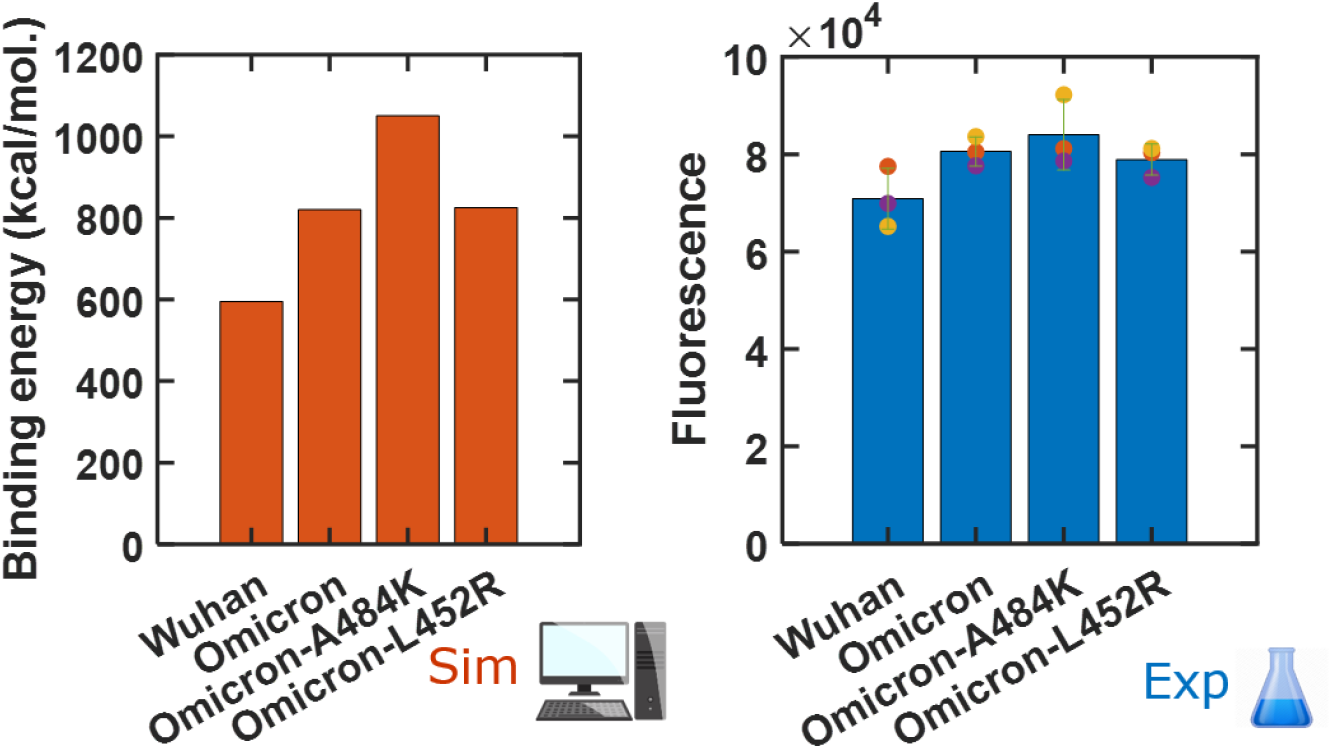
Computational predictions are consistent with the results of binding experiments. Fluorescence results are plotted as the average of MFI across three biological replicates.

## Discussion

In this work, we perform a full QM simulation of intermolecular binding to draw mechanistic insight about the role of each amino acid. We also predict the effect of amino acid mutations on binding at and away from the mutated residue. We show that, in the context of the SARS-CoV-2 S-hACE2 binding, QM-CR can predict the strength of binding when introducing a few mutations into the viral spike by experimentally validating such predictions.

We do not to expect our predictions on binding efficacy to directly correlate to the infectivity of specific SARS-CoV-2 variants. This is well exemplified by the case of mutation L452R, known to improve infectivity via increased fusogenicity [15], a mandatory event for cell entry subsequent of binding. In line with previous research, our simulations do not predict a binding advantage of L452R to hACE2. Our modeling and analysis is focused on the binding effect between the spike and its receptor. Such an analysis does not predict the effects a given mutation may have on other stages of viral infection and replication, especially when conformational changes are implied.

QM-CR relies on a crystal structure as its input, and is therefore dependent on the availability of the crystal structure at an adequate resolution. Such crystal structures ideally include the bound substrate of interest; however, docking simulations can be performed to infer a ligand’s position to inform the model. In our assessment of mutants, we have further assumed that imposing these mutations does not considerably affect the structure; therefore, we have approximated the mutated structure by performing a local optimization around the existing structure [5]. If there are major changes in the shape of a protein as a consequence of mutations, a more representative crystal structure of such variants is needed as the starting point.

In its present stage, QM-CR is a powerful tool if focused on binding energy/affinity. It is therefore of potential use in identifying candidates for molecular inhibitors, antibodies, or any other interactor which is expected to bind and stick to a receptor/substrate. We believe QM-CR can be of particular importance in the context of drug discovery and immunology, especially in the screening for good candidate molecules or for the refinement of existing ones. As a general approach, with little requirement for prior knowledge about interacting molecules, QM-CR can be applied to a wide range of cases when a crystal structure (or a reliable estimate of it) is available. In the case of SARS-CoV-2, QM-CR can inform the design of specific molecular inhibitors against hypothetical variants that may emerge as an important aspect of pandemic readiness.

## Materials and Methods

### Plasmid

The pCAGGS plasmid expressing SARS-CoV-2 Wuhan strain spike protein has previously been described [16]. Synthetic DNA of spike genes corresponding to variants of SARS-CoV-2 omicron strains were obtained from Integrated DNA Technologies (IDT, Coralville, IA, USA). The variants (sequences available in table S1) were cloned into pCAGGS in replacement of SARS-CoV-2 Wuhan strain spike protein via KpnI/SacI digestion and T4 DNA ligation. The cloning strategy results in RBD fusion with human IgG1-Fc to generate recombinant RBD-Fc variants. HEK293T (human embryonic kidney; ATCC) were maintained in growth media DMEM 10x composed of Dulbecco’s Modified Eagle Medium (DMEM, Thermo-Fisher, Massachusetts, USA) supplemented with 2mM Glutamine (Thermo-Fisher, Massachusetts, USA), 1% non-essential amino acids (Thermo-Fisher, Massachusetts, USA), 100 U/mL penicillin and 100 μg/mL streptomycin (Thermo-Fisher, Massachusetts, USA), and 10% FBS (Thermo-Fisher, Massachusetts, USA) at 37°C in 5% CO_2_. HEK293T Cell lines expressing human ACE2 (hACE2) were previously described [16]. Parental cells were transduced with generated MLV virus, and the hACE2 stably expressing cell lines were selected and maintained with medium containing 3 μg/ml puromycin (Sigma, Missouri, USA). hACE2 expression was confirmed by immunofluorescence staining using mouse monoclonal antibody against c-Myc antibody 9E10 (Thermo-Fisher, Massachusetts, USA) and goat-anti-mouse IgG APC (Jackson ImmunoResearch Laboratories, Pennsylvania, USA).

### Protein production and purification

Fc fusion protein production and purification has been previously described RBD-Fc [17,18]. Briefly, HEK293T cells were culture in DMEM 10x media. 12×10^6 293T cells were seeded in T175 flask (ThermoFisher, Massachusetts, USA) and incubated overnight (37ºC, 5% CO_2_) on the day before transfection. On the day of transfection, plasmid DNA were transfected using 70 ul of Transfectin Lipid Reagent (BioRad, California, USA). 4 or 5 T175 plates were prepared for each plasmid DNA. 24 hours after initiation of transfection, culture media was replaced with serum free media (SFM) composed of DMEM 10x composed of Dulbecco’s Modified Eagle Medium (DMEM, Thermo-Fisher, Massachusetts, USA) supplemented with 2mM Glutamine (Thermo-Fisher, Massachusetts, USA), 1% non-essential amino acids (Thermo-Fisher, Massachusetts, USA), 100 U/mL penicillin and 100 μg/mL streptomycin (Thermo-Fisher, Massachusetts, USA). 5 days after transfection, culture supernatants were harvested and cell debris were pelleted by centrifugation (1500 rpm, 5 minutes). The resulting supernatant was incubated overnight with protein A agarose beads at 4ºC with rotation. RBD-Fc then purified by gravity flow using Econo-Glass column (BioRad, California, USA). RBD-Fc elution was performed using 0.1 M glycine (pH2.2) and the eluate was neutralized with 1.5 M Tris-HCl pH 8.8. Purified protein was dialyzed overnight against pH 7.4 phosphate buffer saline (PBS) using 10,000 Molecular Weight Cut-Off (MWCO) dialysis cassettes (BioRad, California, USA). Finally, purified RBD-Fc proteins were concentrated using 10,000 MWCO Amicon centrifugation columns (BioRad, California, USA) and stored at 4°C before use. Protein concentration was determined by spectrometry using a NanoDrop microvolume spectrometer (Thermo-Fisher, Massachusetts, USA).

### Flow cytometry to test the binding of coronavirus RBD-Fc proteins to hACE2 receptor

HEK293T Cell lines expressing human ACE2 (hACE2) were cultured as previously described, in presence of 3 ug/ml puromycin to select for hACE2 expression [16]. hACE2 expression was detected using mouse monoclonal antibody against c-Myc-FITC antibody (clone 9E10) from Thermo-Fisher (Massachusetts, USA). Original HEK293T Cell lines were used as a control. Binding affinity of RBD-Fc variants were detected by staining HEK293T-hACE2 cells as previously described [16]. Briefly, 10^6^ cells were incubated (15 min, room temperature) with 0.5 µg/ml of RBD-Fc variants in PBS containing 0.5% bovine serum albumin (BSA) (Thermo-Fisher, Massachusetts, USA). The cells were washed wish PBS, 0.5% BSA and RBD-Fc binding was detected with goat-anti-human-Ig-APC (Jackson ImmunoResearch Laboratories, Pennsylvania, USA). Data was acquired using by flow cytometry using an LSRII (BD Biosciences, California, USA) and analyzed using FlowJo software (FlowJo, LLC).

### Quantum Mechanical Calculations

We performed Kohn-Sham density functional theory (10.1103/PhysRev.145.561) calculations using the BigDFT program with Hartwigsen-Goedecker-Hutter pseudopotentials (<u>10.1063/1.4793260</u>), a grid spacing of 0.4 Angstroms, and the PBE exchange-correlation functional. Dispersion interactions were added using the D3 correction term (<u>10.1063/1.3382344</u>). In BigDFT, the Kohn-Sham orbitals are expanded in a set of in-situ optimized, localized orbitals which are in turn represented by Daubechies wavelets (<u>10.1063/1.4871876</u>). Calculations were performed in gas phase (which was shown to perform adequately compared to the inclusion of implicit solvent for this system in our previous work [5] at finite electronic temperature using the CheSS library (<u>10.1021/acs.jctc.7b00348</u>).

We postprocess these QM calculations to compute interactions measures between the various components of the systems. We define a general quasi-observable associated with two system fragments:

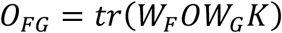

where K is the density matrix of the full system, W the projection on to a given fragment, and O the operator of interest. In this formulation, the fragment bond order is defined as:

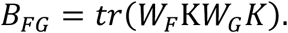

The short/range chemical interaction is defined as the sum of (H the Kohn-Sham Hamiltonian).:

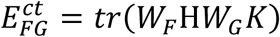

and the empirical D3 correction. The final contribution to the interaction energy is the long/range electrostatic term (ρ the charge density):

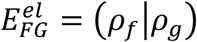

which is approximated as the sum of atom centered multipoles (up to second order) computed from the electronic density.

## Supplementary Materials

**Table S1.**
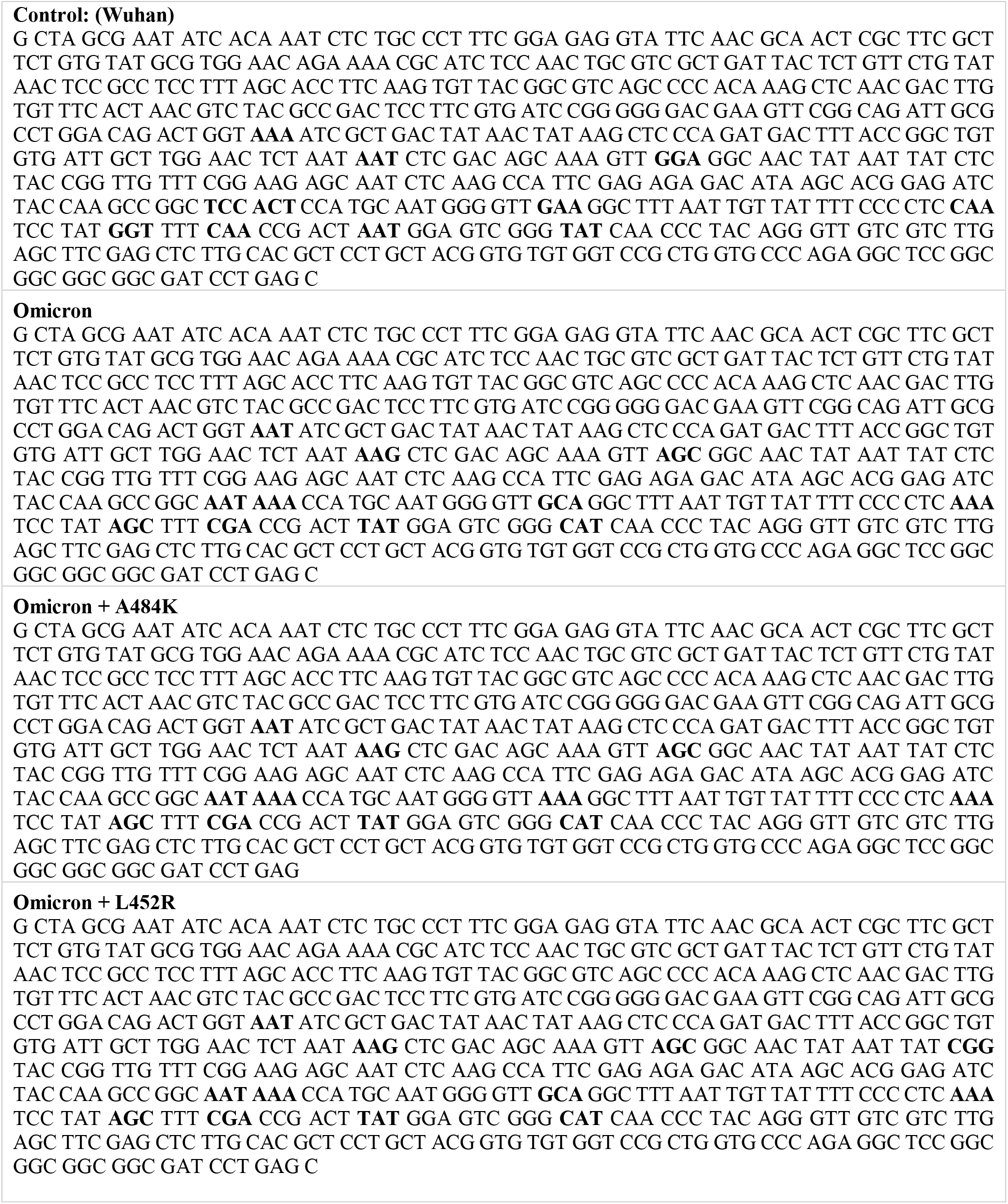
Nucleotide sequences of the various SARS-CoV-2 S constructs. The highlighted codons are those that may differ across mutants and/or compared to the Wuhan reference spike.

**Table S2.** Comparisons between different data obtained from flow cytometry show significance difference between observed binding of hACE2 to different variants of SARS-CoV-2 spike. Calculated *p*-values are obtained from Mann-Whitney U test of APC_A readouts from the flow cytometer using the `ranksum` routine in Matlab.

|  | **WT vs**<br>**Omicron** | **WT vs**<br>**Omicron-A484K** | **WT vs**<br>**Omicron-L452R** | **Omicron vs**<br>**Omicron-L452R** |
| --- | --- | --- | --- | --- |
| *p*-value | $1.0 \times 10^{-20}$ | $6.8 \times 10^{-148}$ | $4.2 \times 10^{-9}$ | $1.6 \times 10^{-3}$ |

**Fig S1.**
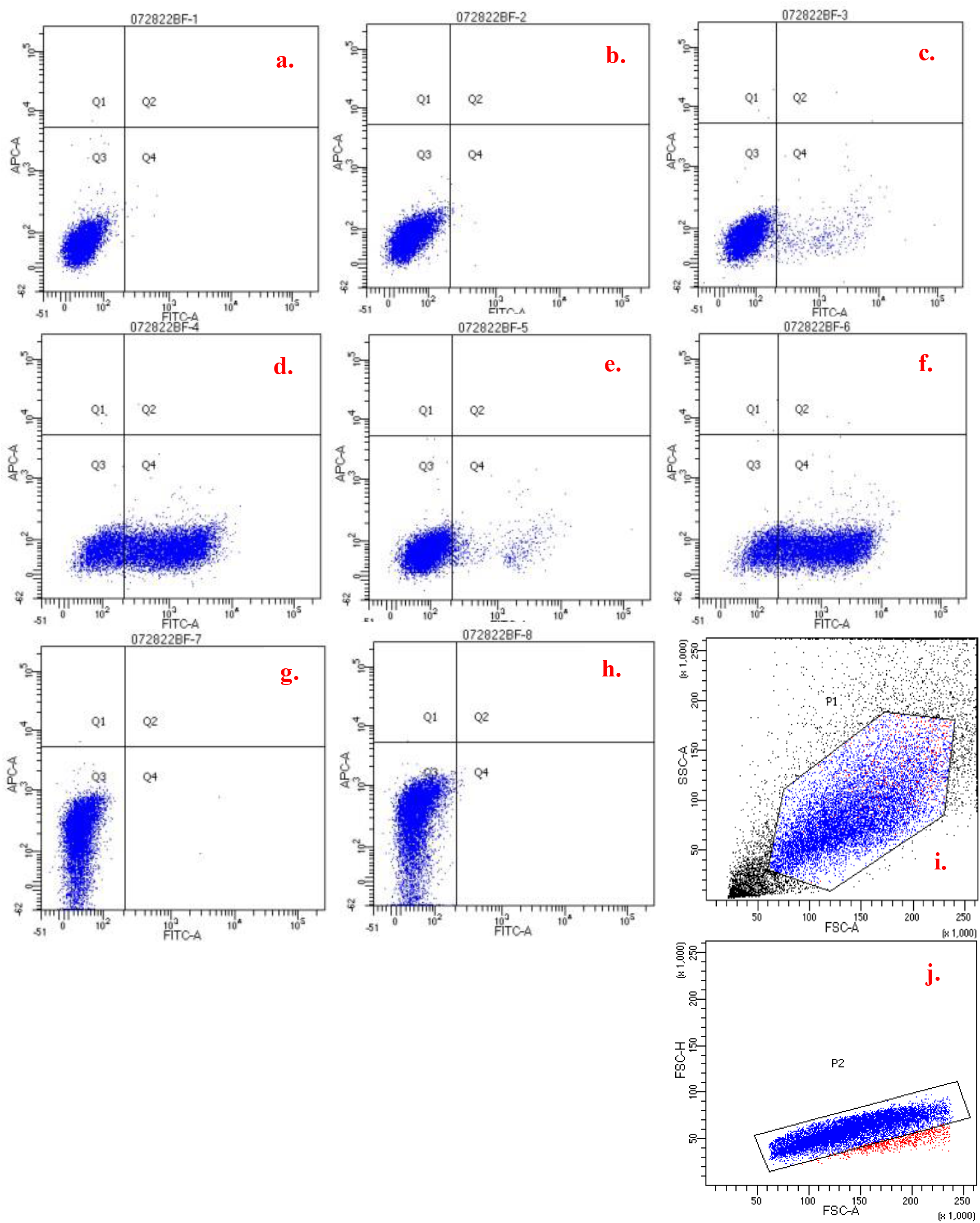
Gating scheme for binding assay. (a) 293T cells; (b) 293T witch ACE2; (c) 293T cells with 3 uL c-myc; (d) 293T cells with ACE2 and 3 uL c-myc; (e) 293T cells witch 5 uL c-myc; (f) 293T cells with ACE2 and 5 uL c-myc; (g) 293T cells with 1 uL hIg-APC; (h) 293T cells with ACE2 and 1 uL hIg-APC; (i) gating for live cells; (j) gating for single cells.

**Fig S2.**
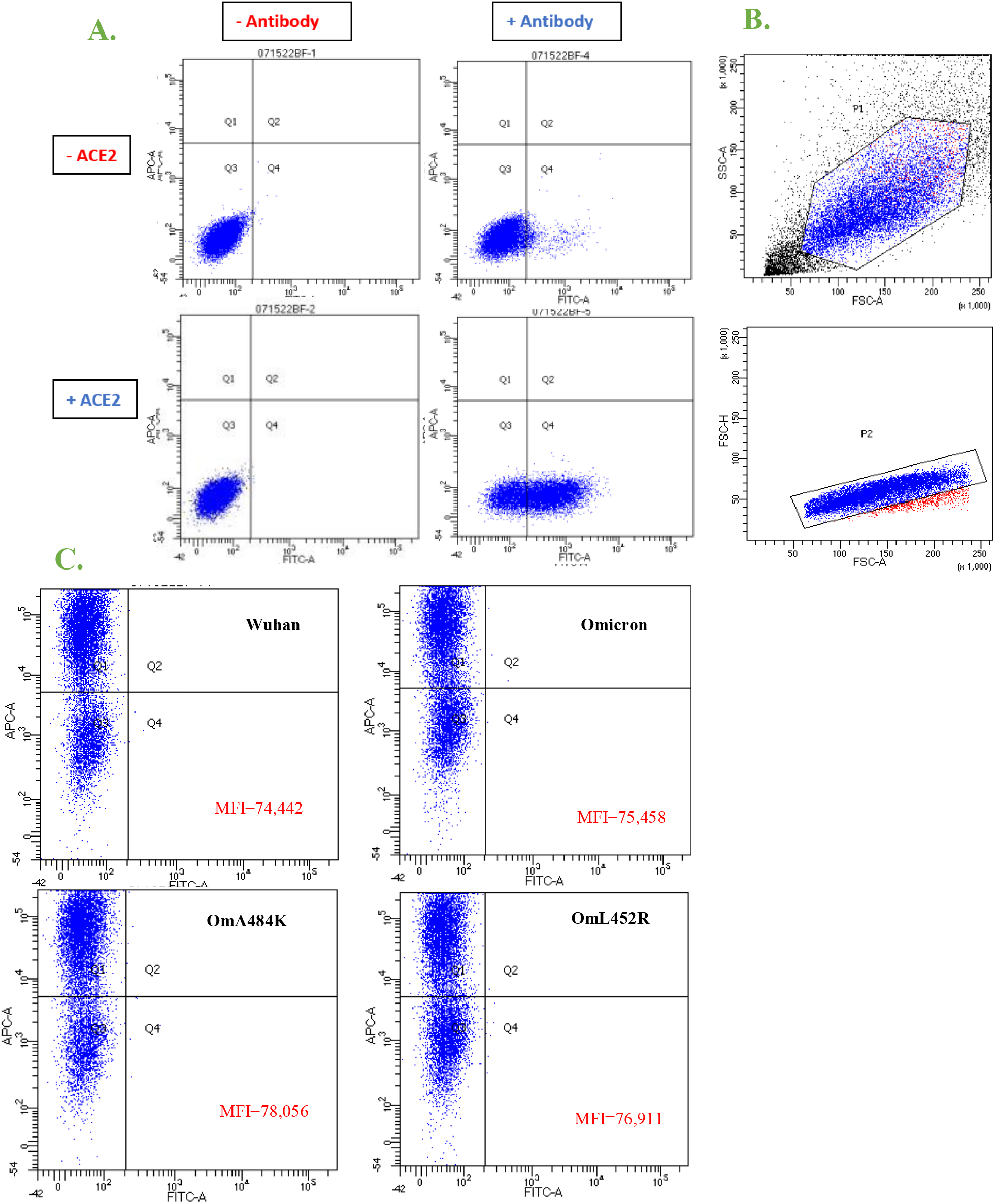
First replicate of binding experiment. Panel A shows fluorescence across all combinations of stained and non-stained cells, with and without ACE2 expression. Panel B shows gating on live (a) and single cells (b). In panel C, the four different S protein binding performance are expressed in mean fluorescence intensity (MFI) after APC-A binding,

**Fig S3.**
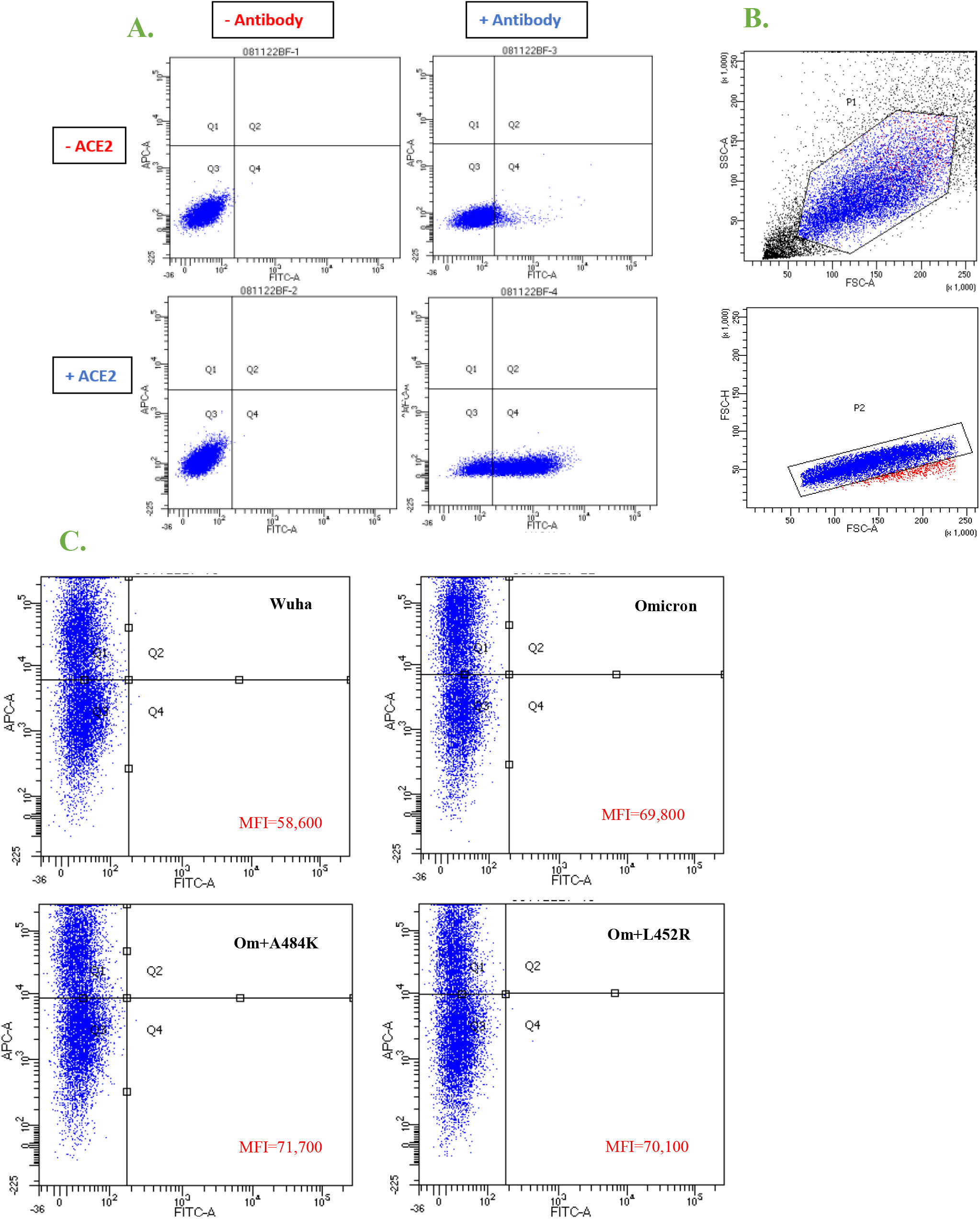
Second replicate of binding experiment. Panel A shows fluorescence across all combinations of stained and non-stained cells, with and without ACE2 expression. Panel B shows gating on live (a) and single cells (b). In panel C, the four different S protein binding performance are expressed in mean fluorescence intensity (MFI) after APC-A binding.

**Fig S4.**
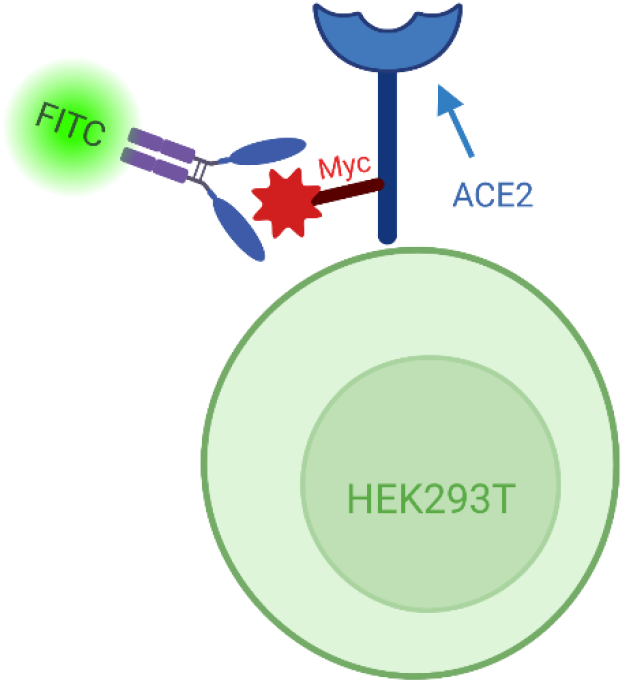
Visual scheme of the fluorimetric assay. To verify 293T cells expression of ACE2, the myc tag on ACE2 is bound with an antibody carrying the FITC fluorescent label for detection.

**Fig S5.**
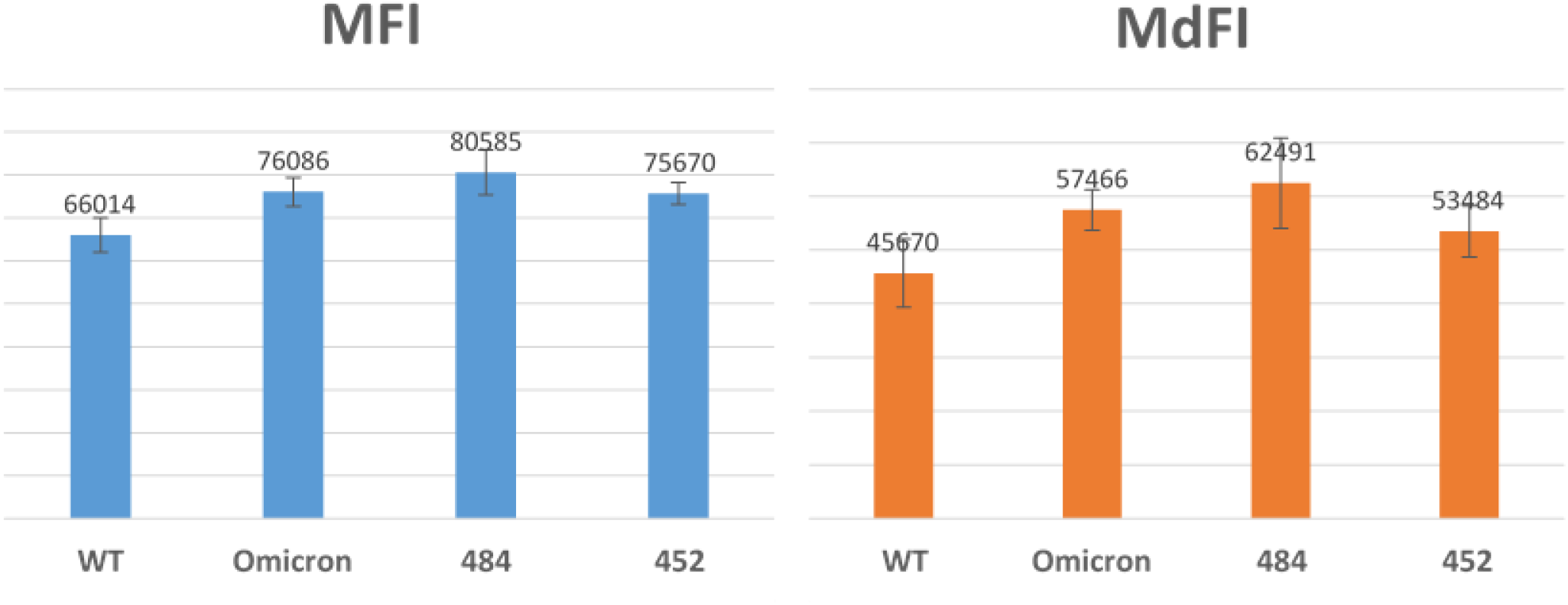
Overall mean and median fluorescence intensities show similar trends. Mean (MFI) and median (MdFI) fluorescence of APC-A readouts are compiled across all the three experiments performed.

## Author contributions

Conceptualization: MZ, LG, MF, BM

Methodology: MZ, LG, WD, IF, BL, PA, AJ, HJ

Investigation: MZ, LG

Visualization: MZ, LG, WD, IF, BM, PA

Funding acquisition: MF, BM, LG, WJ, IF, TN

Project administration: BM, LG

Supervision: MF, BM

Writing – original draft: MZ, LG, BM

Writing – review & editing: MZ, LG, BM, BL, IF, WJ, MF

## Data and materials availability

All data, code, and materials used in the analysis must be available in some form to any researcher for purposes of reproducing or extending the analysis. Include a note explaining any restrictions on materials, such as materials transfer agreements (MTAs). Note accession numbers to any data relating to the paper and deposited in a public database; include a brief description of the data set or model with the number. If all data are in the paper and supplementary materials, include the sentence “All data are available in the main text or the supplementary materials.”

## Acknowledgements

We acknowledge useful discussions and support by Michel Masella, Lorenzo Fontolan, Massimo Reverberi, Andrea Kirmaier, Huihui Mou, Michael Travisano. We thank Patrick Autissier and the Boston College Flow Cytometry Core for infrastructure and support. Work in the Momeni lab was supported by a by an NSF-CBET Env. Engineering award (NSF#2103545). LG acknowledges support from the European Centre of Excellence MaX (project ID 676598). Illustrations in figures 3 and S4 were created in Biorender.com.

